# A DNA nanoassembly-based approach to map membrane protein nanoenvironments

**DOI:** 10.1101/836049

**Authors:** Elena Ambrosetti, Giulio Bernardinelli, Ian Hoffecker, Leonard Hartmanis, Rickard Sandberg, Björn Högberg, Ana I. Teixeira

**Affiliations:** Department of Medical Biochemistry and Biophysics, Karolinska Institutet, Sweden; Department of Cell and Molecular Biology, Karolinska Institutet, Sweden

## Abstract

Super-resolution imaging has revealed that most proteins at the plasma membrane are not uniformly distributed but localize to dynamic domains of nanoscale dimensions. To investigate their functional relevance, there is a need for methods that enable comprehensive mapping of the compositions and spatial organizations of membrane protein nanodomains in cell populations. However, current superresolution methods are limited to analysing small, preselected subsets of proteins, at very low sampling fractions. Here we describe the development of a non-microscopy based super-resolution method for unbiased ensemble analysis of membrane protein nanodomains. The method, termed NANOscale DEciphEring of membrane Protein nanodomains (NanoDeep), is based on the use of DNA nanoassemblies to translate membrane protein organization information into a DNA sequencing readout. Using NanoDeep, we characterized the nanoenvironments of Her2, a membrane receptor of critical relevance in cancer. We found that the occupancies of Her2, Her3 and EGFR in the nanoenvironments surrounding Her2 were similar in two cell lines with vastly different expression levels of Her2. Further, we found that adding Heregulin-β1 to cancer cells led to increased occupancy of Her2 and Her3, and to a lesser extent EGFR, in Her2 nanoenvironments. NanoDeep has the potential to provide new insights into the roles of the composition and spatial organization of protein nanoenvironments in the regulation of membrane protein function.

## Main

Cells sense extracellular signals, such as protein ligands, through specialized proteins present on the cell surface called membrane receptors. The protein nanoenvironment, i.e. the composition and spatial organization of proteins surrounding membrane receptors, is dynamic and often modulated by ligand binding, suggesting that it has functional relevance^1, 2^. Super-resolution microscopy has enabled the characterization of the nanoscale spatial distributions of proteins at the cell membrane but is limited to simultaneously analysing only a few proteins. Further, using super-resolution microscopy, it is only feasible to image a small fraction of all protein nanodomains present in a cell population^3–5^. DNA detection as a proxy for protein detection through the use of oligo-conjugated affinity binders has been used extensively for signal amplification^6^ and analysis of proximity between pairs of proteins^7, 8^. Further, DNA sequencing is used as a readout in DNA microscopy, a new method to visualize the spatial organization of RNA and DNA molecules inside cells^9–11^. Here we present NanoDeep, a method that uses DNA sequencing to decipher the nanoscale spatial distribution of membrane proteins. This method allows for the detection *en masse* of the inventory of proteins that forms the nanoenvironment of any reference membrane protein in cell populations.

We demonstrate the application of NanoDeep to the analysis of protein nanoenviroments surrounding Human Epidermal Growth Factor Receptor 2 (Her2). Her2 cooperates with members of the Epithelial Growth Factor Receptor (EGFR) family of proteins (EGFR, Her2, Her3 and Her4) to regulate cell proliferation and differentiation during normal embryonic development^12^. In several cancers (including breast, ovarian and gastric cancers) Her2 is overexpressed and its expression levels correlate with poor prognosis^12, 13^. Interestingly, the oncogenic capacity of Her2 is closely connected to the impact of overexpression on the frequency distributions of interactions between Her2 and EGFR family members at the cell membrane^14–16^. For example, Her2 overexpression leads to increased levels of Her2 and Her3 heterodimers, which drive more potent oncogenic signalling activity than the corresponding homodimers^13, 17–19^. Further, Her2-EGFR dimerization, driven by overexpression of one or both proteins, has been shown to lead to a more aggressive breast cancer phenotype^20^. Notably, new evidence supports the hypothesis that not only the formation of dimers but also of higher-order receptor assemblies at cell membrane regulates Her2 function^21–25^. Although there are well established correlations between the levels of Her2 homo- and heterodimers and cancer aggressiveness, the roles of the composition of Her2 protein nanoenvironments for downstream signalling are poorly understood^19^. Using NanoDeep, we characterized the protein nanoenvironments of Her2 in model surfaces and in cells. We found that SKBR3 breast cancer cells that overexpress Her2 showed similar levels of occupancy of Her2, Her3 and EGFR in Her2 nanoenvironments compared to MCF7 breast cancer cells, which present basal levels of Her2. However, the higher expression levels of Her2 in SKBR3 cells correlated with higher total levels of Her2, Her3 and EGFR in Her2 nanoenvironments, compared to MCF7 cells. Further, stimulation of SKBR3 cells with the Her3 ligand Heregulin-β1 (HRG-β1) led to an increase in occupancy of Her2 and Her3, and to a lesser extent of EGFR, in Her2 nanoevinronments. Together, these results indicate that NanoDeep is able to characterize differences in protein nanoenvironments in different cellular contexts.

### NanoDeep

The NanoDeep method converts protein spatial distribution information into a DNA sequencing readout (Fig. 1). We designed a DNA nanoassembly, which we named NanoComb, composed of four single-stranded DNA (ssDNA) oligos, called prongs, that protrude from a doublestranded backbone at regular intervals. The prongs contain a barcode that identifies their position within the NanoComb. The first prong is defined as the reference prong and the remaining prongs are the detection prongs. The reference prong is preloaded with a binder that recognizes a reference protein, which is Her2 in our model workflow. This is done by conjugating the binder for the reference protein with an oligo that partially hybridizes with the reference prong (Fig. 1a). After incubating the NanoComb bearing the binder for the reference protein with fixed cells and washing, a library of binders for the inventory of proteins to be analysed is added, each conjugated to an oligo containing a barcode that identifies the protein recognized by the binder (Fig. 1b). The oligos further contain a sequence, which is common to all binder-oligo conjugates, that is partially complementary to the detection prongs. Importantly, to prevent binder-oligo conjugates that are not bound to membrane proteins from hybridizing with the detection prongs, hybridization is blocked when conjugates are added to the cells and unblocked after washing away conjugates that are not bound to their target membrane proteins (Fig. 1c). The hybridization between the prongs and the binder oligos creates free 3’ ends that act as primers for DNA polymerase (Fig. 1d), leading to the formation of double-stranded DNA (dsDNA) sequences that contain both the barcode for the position of the prong as well as the binder barcode (Fig. 1e). As specific nuclease sequences are incorporated in both the prongs and binder oligos, these dsDNA sequences can be cleaved by a nuclease (Fig. 1f) and analysed by next generation sequencing (NGS) (Fig. 1g), providing information on the composition and spatial organization of the nanoenvironment surrounding the reference protein across the cell population.

**Fig. 1:**
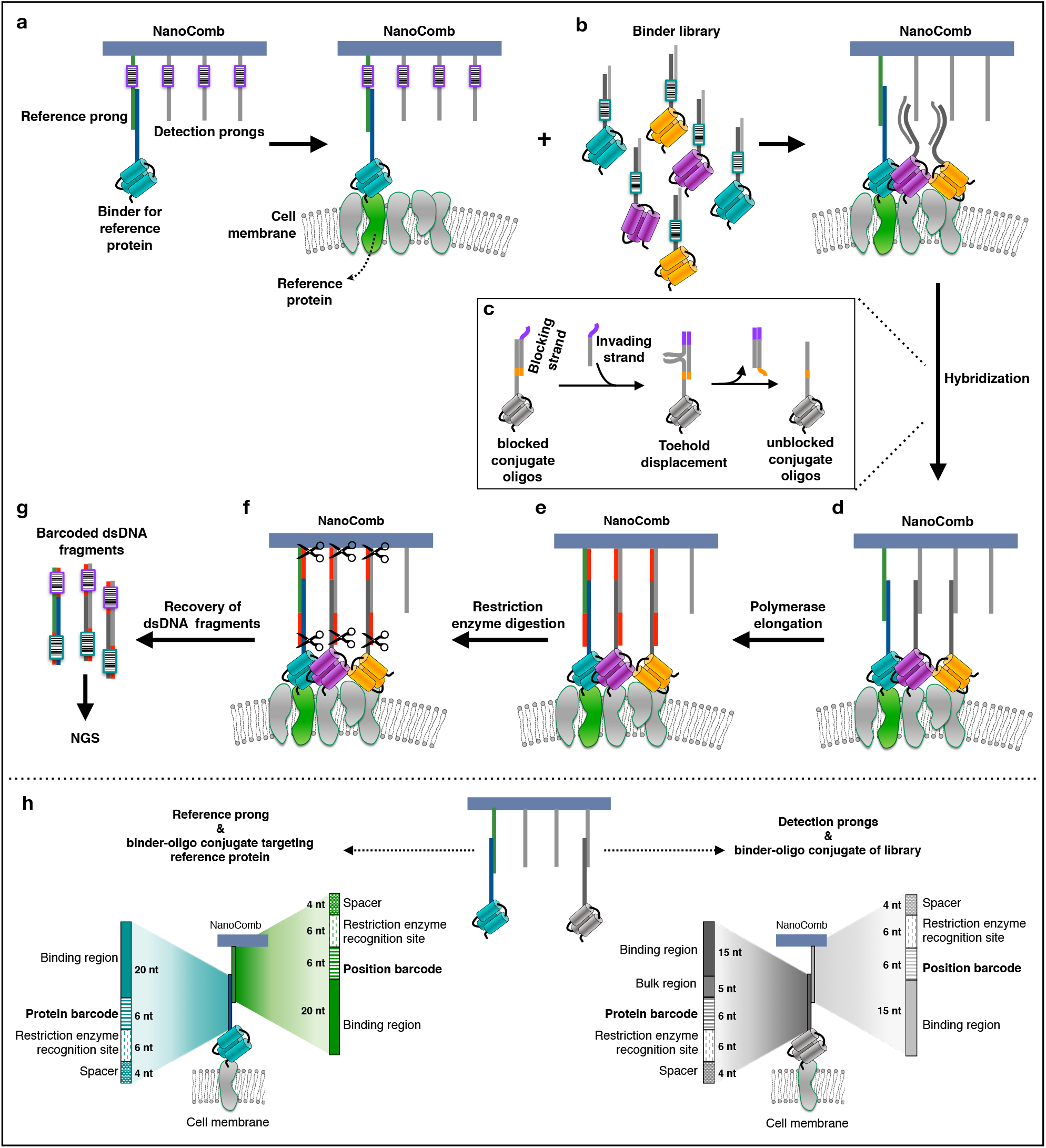
Schematic of the NanoDeep method. **a**, A DNA nanoassembly (NanoComb) consisting of a double stranded backbone with four barcoded protruding ssDNA strands (prongs) is preloaded with an oligo-conjugated binder specific for the reference protein and incubated with fixed cells. **b**, A library of binders is added, each conjugated to a ssDNA sequence bearing a barcode that identifies its target protein as well as a sequence that is partially complementary to the sequences of the detection prongs. **c**, During the incubation with the target proteins, affibody-oligo conjugates are hybridized with a blocking strand to provide a blocked configuration. Unblocking is promoted by toehold-mediated displacement guided by an invading strand. **d**, The free 3’ ends formed by the hybridization of the prongs with the binder oligos function as primers for DNA polymerase. **e**, DNA polymerase reaction creates dsDNA sequences that contain both the barcodes for the relative position of the prongs within the NanoComb and for the protein that is recognized by the binder. **f**, Restriction enzymes cleave the dsDNA sequences at specific nuclease target sequences that are included in both the prongs and the binder oligos, leading to the release of dsDNA sequences. **g**, dsDNA sequences are analysed by Next Generation Sequencing. **h**, Schematic representation of prongs and oligos conjugated to the binders, which contain binding regions complementary to each other. For the reference prong and the oligo conjugated to the binder targeting the reference protein (left) this region is 20-nucleotides (nt) long. For detection prongs and the oligos conjugated to the library binders (right) the binding region is 15-nt long; a 5-nt long bulk region is added to the oligo sequences of the conjugates. Binding regions are followed by a 6-nt barcode identifying the protein or the position, in the binder oligos or the prongs, respectively. Further, both the prongs and the binder oligos contain nuclease target sites (6-nt) followed by a 4-nt spacer, included to facilitate the binding of the restriction enzyme.

Super-resolution microscopy studies determined that the spacing of proteins in Her2 dimers is on average in the range of 10-20 nm^15, 23, 26^ and that Her2 clusters in breast carcinoma cell lines have a mean diameter of 67 nm^21^. The detection prongs of the NanoCombs are positioned by design at 7, 14 and 21 nm relative to the reference prong, in an extended DNA conformation. Therefore, the geometry of NanoCombs enables probing of the relevant length scale for the analysis of Her2 nanoenvironments.

### Design, production and characterization of DNA NanoCombs and binder-oligo conjugates

NanoCombs were produced by hybridizing a 100-nucleotide (nt) long ssDNA oligo (backbone) to four shorter oligos (prongs) partially complementary to the backbone (Fig. 2a,b). The prongs form a pattern of four ssDNA oligos with a period of 21 base pairs (bp), protruding from the same side of the backbone. The period of the prongs corresponds to a distance of 7 nm in an extended DNA conformation. The length of the protruding portion of the prongs is 36 and 31 bp for reference prong and detection prongs, respectively, corresponding to a maximum length of 12.5 nm, when doublestranded (Fig. 2a,b).

**Fig. 2:**
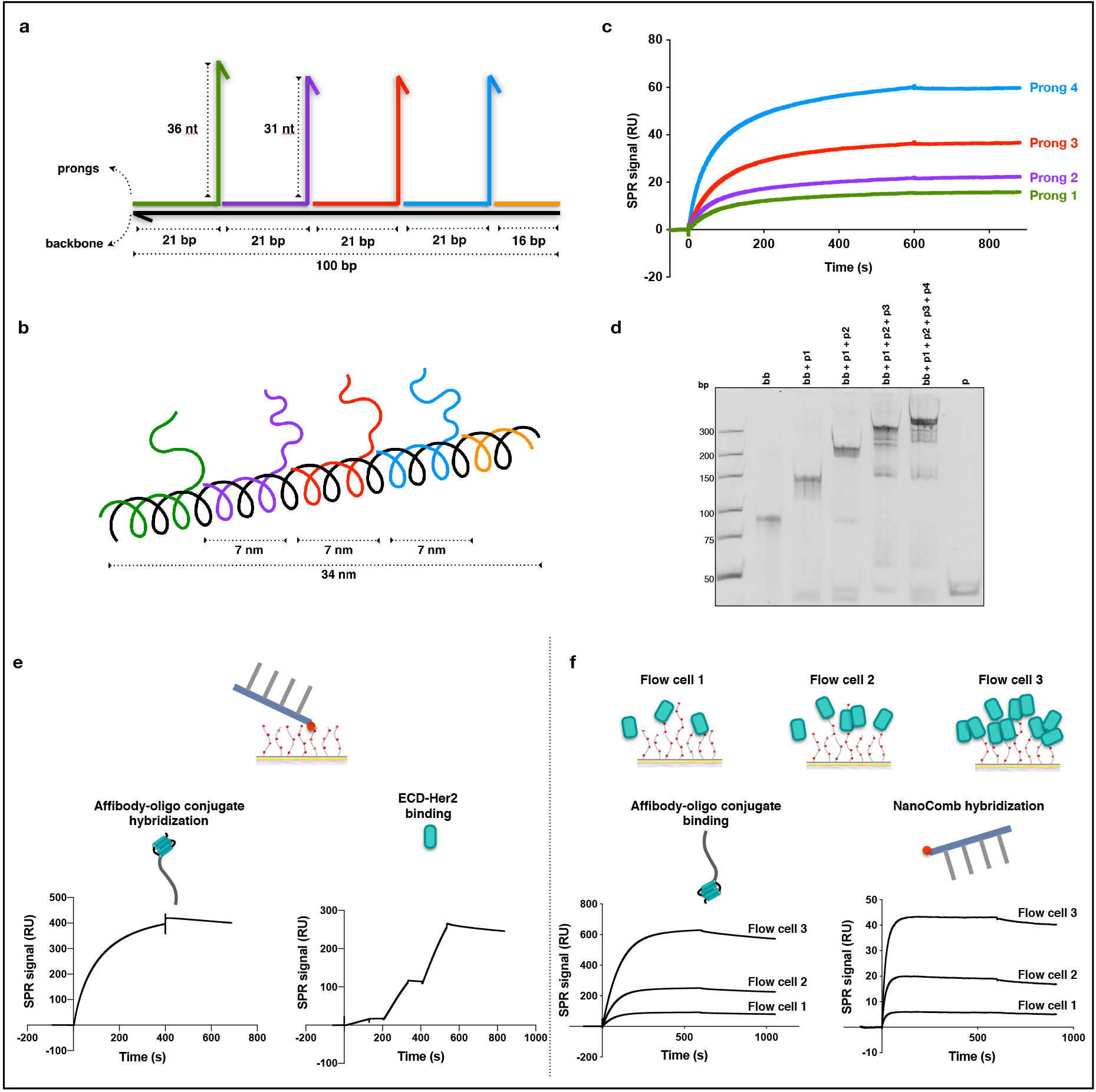
NanoComb characterization. **a**, NanoComb composed of a 100-nt ssDNA backbone and four 57- or 52-nt ssDNA strands (reference and detection prongs, respectively), each of which hybridizes to the backbone by means of the first 21-nt, leaving the remaining 36- or 31-nt ssDNA sequences to protrude from the backbone. **b**, In an extended DNA conformation, the total length of NanoComb is 34 nm and the prongs protrude from the backbone with a period of 21 bp, which corresponds to a distance of 7 nm, along the helical direction of the backbone. **c**, Real time kinetic analysis using SPR of sequential binding of the prongs to the backbone, which was immobilized onto the SPR sensor surface. RU, resonance units. **d**, The backbone alone (bb) or the backbone hybridized with one (bb+p1), two (bb+p1+p2), three (bb+p1+p2+p3) or all four prongs (bb+p1+p2+p3+p4), monitored by native PAGE (7%). One prong sequence was loaded as a control (p). **e, f**, The incorporation of affibody-oligo conjugates in the NanoCombs was verified using direct (e) and reverse (f) SPR assays. In the direct assay (e), we measured the hybridization of affibody-oligo conjugates to desthiobiotinylated NanoCombs immobilized on Streptavidin (SA) SPR surfaces, followed by measurement of ECD-Her2 binding, to verify that the affibodies preserved their ability to recognize Her2. In the reverse assay (f), ECD-Her2 was covalently immobilized on the sensor surface at three different surface densities. Sensorgrams showed sequential binding of affibody-oligo conjugates and of NanoCombs. The SPR signal obtained from NanoComb binding to the affibody-oligo conjugates was proportional to the amount of bound conjugates, which in turn reflected the surface density of ECD-Her2.

After folding the NanoCombs, we removed the excess prongs by using streptavidin-coated magnetic beads and taking advantage of the lower affinity of desthiobiotin compared to biotin (Fig. S1). We monitored the assembly of NanoCombs with Surface Plasmon Resonance (SPR) assay, showing sequential hybridization of the four prongs with the backbone (Fig. 2c). We further confirmed NanoComb assembly and purification with native polyacrylamide gel electrophoresis (PAGE), which showed that the four prongs hybridized with the backbone (Fig. 2d) and that the NanoCombs were purified from excess prongs (Fig. S1).

We selected affibodies as binders due to their high affinity and small dimensions, minimizing the impact of the size of the binder *per se* on the spatial resolution of the method. To enable site specific and stoichiometric conjugation of affibodies to DNA oligos, we used a conjugation approach based on a self-labelling tag derived from a truncated VirD2 protein of *Agrobacterium tumefaciens*, that enables fast conjugation of unmodified DNA molecules to proteins in physiological conditions and with a controlled stoichiometry of 1:1^*27*^. We produced fusion proteins between VirD2 and affibodies that bind the extracellular domains of three members of the EGFR family (EGFR, Her2 and Her3). SPR assay showed that all affibodies fused to VirD2 exhibited high affinity and selectivity for their specific targets (Fig. S2). Further, native PAGE demonstrated efficient conjugation of the affibodies with the oligos (Fig. S3a). We observed that the oligo conjugation caused a slight increase in the KD of the affibodies that nevertheless did not prevent them from recognizing their targets with high affinity since the KD was still in the nanomolar range, and, more significantly, the low dissociation rate (koff) was preserved (Fig. S3b).

To assess the hybridization of the affibody-oligo conjugates to the NanoCombs, we used direct and reverse SPR assays using anti-Her2 affibody-oligo conjugates as a test sample. We observed that anti-Her2 affibody-oligo conjugates hybridized effectively with the NanoCombs and that the capability to recognize the specific target was preserved (Fig. 2e). Further, NanoComb binding was proportional to the levels of affibody-oligo conjugates bound to the SPR surface (Fig. 2f). Together, these results support the specificity of the interaction between the NanoCombs and the affibody-oligo conjugates.

### Toehold exchange strategy to reversibly block the hybridization of affibody-oligo conjugates to the detection prongs of the NanoCombs

The NanoDeep method consists of first targeting a reference protein and then detecting its protein nanoenvironment. In the model workflow presented here, the targeting step consists of binding NanoCombs preloaded with anti-Her2 affibody (Her2-NanoCombs) to Her2. The detection step involves binding of anti-Her2, -Her3 and -EGFR affibody-oligo conjugates (binder library) to their targets and hybridization of the oligos to the detection prongs of the NanoCombs. Therefore, it is crucial that the hybridization of the binder library to the detection prongs of the NanoCombs occurs only when the affibodies are first bound to their target proteins. To address this, we developed a strategy based on toehold-mediated strand displacement^28, 29^, widely used in dynamic DNA nanotechnology. To validate this strategy, biotinylated versions of the binder oligos were anchored to streptavidin coated SPR surfaces (Fig. S4). Hybridization to a blocking strand caused an increase in the SPR signal, which was followed by a decrease in the signal due to displacement of the blocking strand through strand migration upon adding an invading strand. The resulting unblocked oligos were able to hybridize to the NanoCombs, which led to an increase in the SPR signal (Fig. S4a). In the absence of strand invasion, the blocked oligos were not able to bind to the NanoCombs (Fig. S4b). To verify that the affibody-oligo conjugates were able to hybridize, following toehold exchange, to the detection prongs of Her2-NanoCombs, we performed SPR assays using surfaces that presented both ECD-Her2 and ECD-Her3. Her2-NanoCombs were incubated with the functionalised SPR surfaces (Fig. 3). Following washing, anti-Her3 affibody-oligo conjugates that were pre-hybridized to blocking strands were injected. After allowing for binding of the conjugates to ECD-Her3 on the SPR surfaces and washing away unbound conjugates, the blocking strand was displaced by adding the invading strand. Importantly, we performed this step after increasing the temperature to 45 °C, which is below the melting temperature of the hybridization between the invading strand and the blocking strand but above the melting temperature of the hybridization between the invading strand and the prongs. This prevented the hybridization of invading strand to the prongs of the NanoCombs. Temperature was then decreased to 25 °C and the oligo tails of the affibody-oligo conjugates were then free to hybridize to the detection prongs of the NanoCombs. To confirm that the detection prongs of the NanoCombs were able to bind to the unblocked anti-Her3 affibody-oligo conjugates, we introduced a second toehold-mediated exchange system that displaced the hybridization between the reference prong and the anti-Her2 affibody-oligo conjugate. In this manner, the NanoCombs were able to stay bound to the surface only if the anti-Her3 affibody-oligo conjugates were properly unblocked and were able to hybridize to the detection prongs. Accordingly, we did not observe significant loss of the SPR signal upon injection of the invading strand for the anti-Her2 affibody-oligo conjugate on samples where we performed unblocking of the anti-Her3 affibody-oligo conjugates (Fig. 3a-left). In contrast, the SPR signal decreased on surfaces that had not been treated with anti-Her3 conjugates (Fig. 3a-right). Using Electrophoretic Mobility Shift assay (EMSA), we further confirmed that the affibody-oligo conjugates were able to hybridize to the blocking strand and that adding the invading strand reversed this hybridization, when performing the reactions in solution (Fig. 3b). Together, these results show that toehold-mediated strand displacement reversibly blocked the hybridization of affibody-oligo conjugates to the detection prongs of the NanoCombs with high efficiency.

**Fig. 3:**
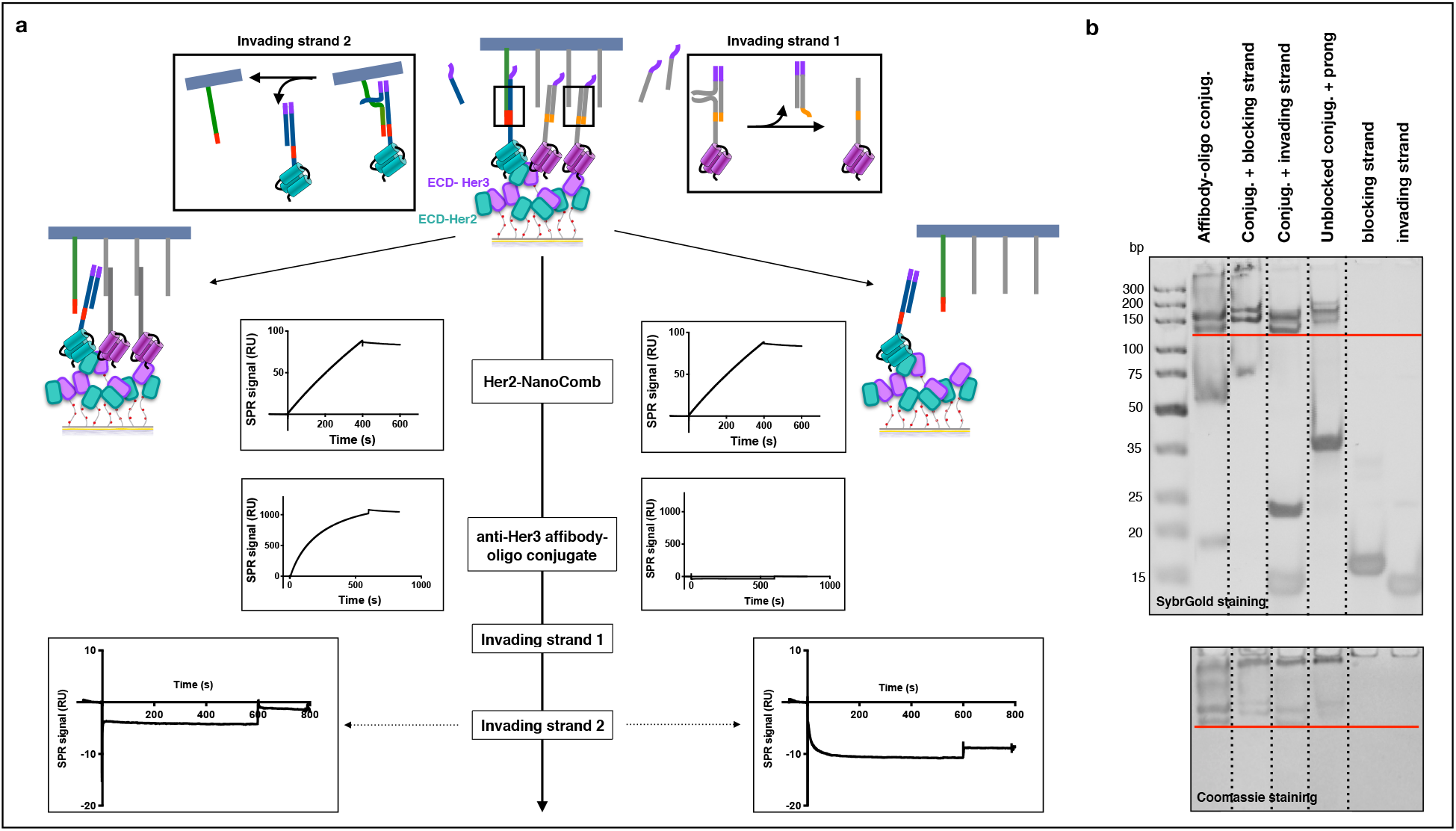
Toehold exchange reversibly blocked the hybridization of affibody-oligo conjugates to the NanoCombs. **a**, A 1:1 mixture of ECD-Her2 and ECD-Her3 was covalently attached to the SPR surfaces of two separate flow cells. Her2-NanoCombs were injected and binding to the anchored ECD-Her2 was detected by an increase in the sensorgram signal. Anti-Her3 affibody-oligo conjugates, previously hybridized with a blocking strand, were added to flow cell 1 (on the left) but not to flow cell 2 (on the right). In flow cell 1, a first invading strand (invading strand 1) was injected, promoting unblocking of the anti-Her3 affibody-oligo conjugates and hybridization to the detection prongs of the NanoComb. The sequences of the anti-Her2 conjugates preloaded on the reference prong contained an added 7-nt at 3’ end in this experiment. This allowed for a second toehold exchange reaction that promoted the displacement of anti-Her2 conjugates from the reference prong. This caused a larger decrease in the SPR signal in flow cell 2, where binding of the NanoComb to the surface is mediated only by the reference prong (bottom-right), than in cell flow 1, where the NanoComb remains bound to the surface through the interaction of the detection prongs with the anti-Her3 conjugates (bottom-left). **b**, EMSA using native PAGE (13%) was performed on samples that underwent blocking/unblocking reactions in solution. The hybridization of blocking strand to affibody-oligo conjugate was visualized by a shift of the conjugate band (the red line indicates the level without blocking). Displacement of the blocking strand by the invading strand was visualised by a shift in the conjugate band and the presence of the released dsDNA fragment consisting of the blocking strand hybridized to the invading strand. We observed a further shift in the conjugate band when adding the complementary prong.

### Validation of enzymatic reactions and recovery of the barcoded dsDNA sequences

Once correct binding is established between the prongs of the NanoCombs and the binder oligos, two enzymatic reactions, DNA polymerization and cleavage, are needed to obtain the final dsDNA sequences that hold both the position and identity barcodes. Using SPR, we were able to monitor the release of NanoCombs and barcoded dsDNA fragments from the surface, following successful T4 polymerase elongation and BamHI/EcoRI cleavage reactions (Fig. 4a,b). We further confirmed with native PAGE that we obtained the final barcoded fragments only from samples where both enzymatic reactions had occurred (Fig. 4c).

**Fig. 4:**
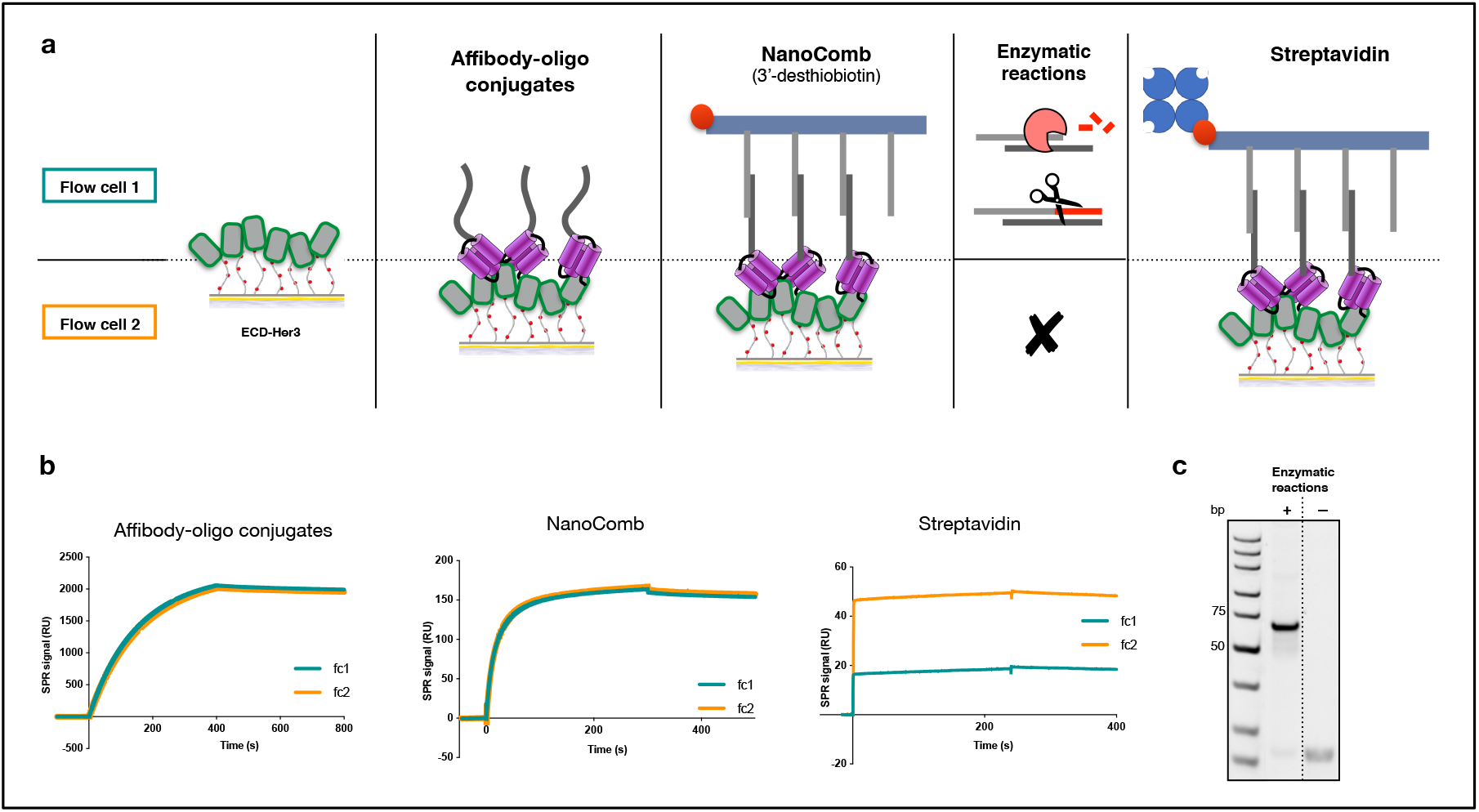
DNA polymerase and nuclease reactions generated barcoded dsDNA sequences. **a**, Schematic representation of the SPR assay; equivalent amounts of E CD-Her3 were covalently attached to two SPR flow cells. Anti-Her3 affibody-oligo conjugates bound first to the target proteins and then hybridized with the NanoCombs. To promote the release of the barcoded dsDNA sequences, T4 polymerase DNA elongation and restriction enzyme cleavage were performed. The enzymatic reactions were carried out only on flow cell 1 and flow cell 2 was used as a negative control. Finally, streptavidin that is able to bind to the desthiobiotin on 3’ end of NanoComb backbone, was injected over the two flow cells to detect the residual amounts of NanoCombs remaining on the surface. **b**, SPR signals of the conjugates and NanoCombs were comparable on the two flow cells. The binding of streptavidin was significantly reduced in flow cell 1 compared to flow cell 2, demonstrating the efficiency of the enzymatic reactions. **c**, Barcoded dsDNA sequences visualized on native PAGE (13%) after PCR amplification were recovered from flow cell 1 but not from the negative control, flow cell 2.

### NGS decoding of dsDNA sequences containing position and protein identity barcodes

We performed NanoDeep on model SPR surfaces, in which we were able to tune the composition and the surface density of bound proteins. We produced SPR surfaces that presented ECD-Her2 and ECD-Her3 at two different surface densities (Fig. 5a-left). We treated the surfaces with Her2-NanoCombs and performed NanoDeep using binder libraries consisting of anti-Her2 and anti-Her3 conjugates. To investigate whether the NGS analysis reflected the density of the proteins present on the surface, we correlated the SPR binding signal of the library (Fig. 5a-center) with the NGS reads originating from the detection prongs (detection sequences), scaled to the reads from the reference prongs (reference sequences). We observed that there was a correspondence between the SPR signal and the number of reads from the detection prongs (Fig. 5a-right). Next, we created surfaces presenting different combinations of the ECD-Her2, -Her3 and -EGFR (Fig. 5b-left) and performed NanoDeep using Her2-NanoCombs and binder libraries consisting of anti-Her2, -Her3 and -EGFR conjugates. NGS analysis of the resulting barcoded dsDNA fragments showed that the detection prongs were able to selectively detect the proteins present in each surface. Importantly, the distribution of reads for position barcodes indicated unbiased probing by the detection prongs of the NanoComb.

**Fig. 5:**
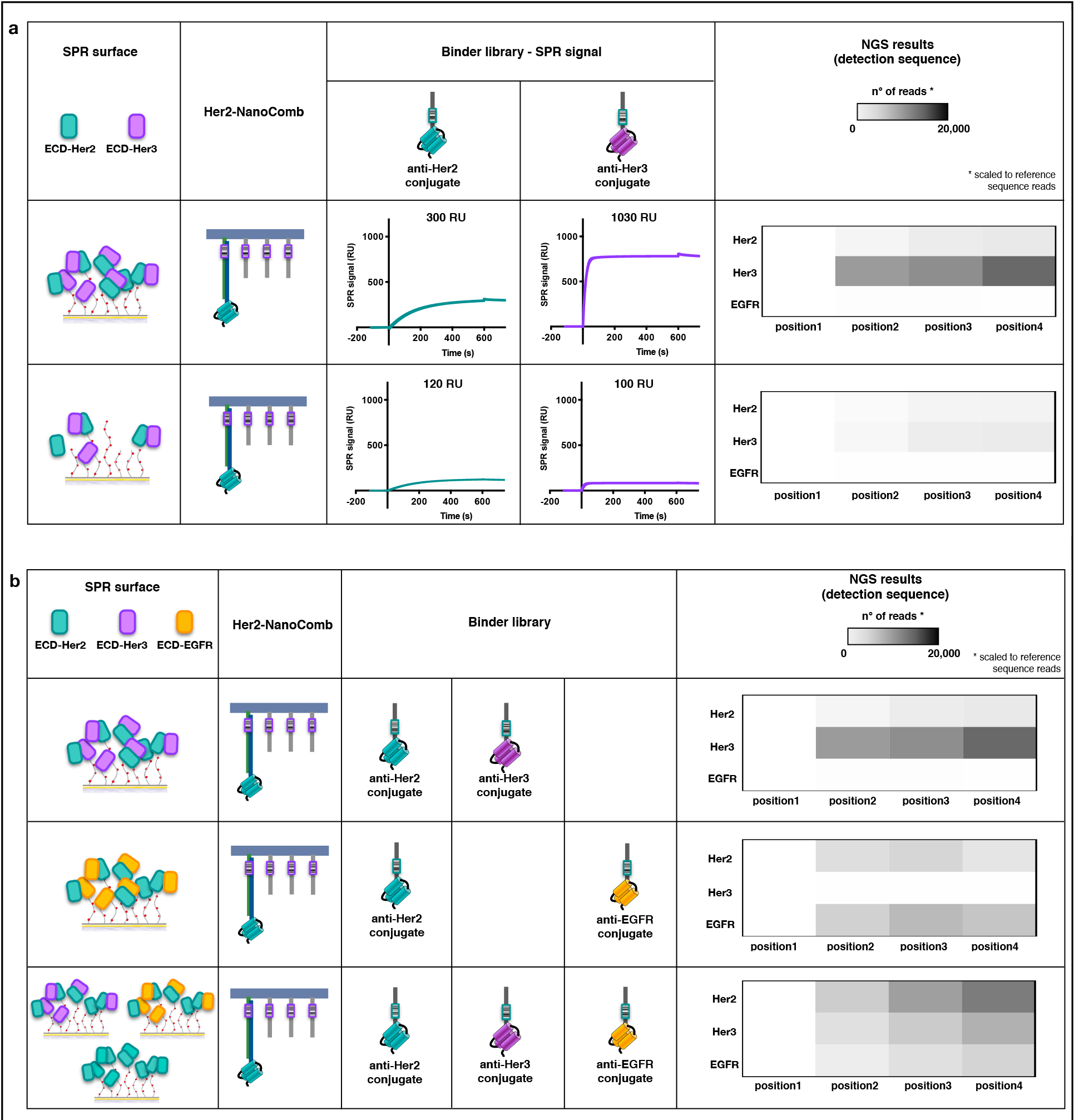
NanoDeep on model SPR surfaces. **a**, 1:1 mixtures of ECD-Her2 and ECD-Her3 were covalently attached to the SPR surfaces of two sensor chips at two different surface densities. NanoDeep was performed by first flowing Her2-NanoCombs and then binder libraries consisting of anti-Her2 and anti-Her3 conjugates. Binding to the anchored proteins was monitored by sensorgram signals, which reflect the amount of anchored proteins. NGS analyses were performed in duplicate and presented as mean values in the heatmap. Barcode reads from the detection sequences were scaled to the reference sequence reads. **b**, NanoDeep was performed on SPR surfaces presenting different compositions of EGFR family receptors: Her2-Her3 (top), Her2-EGFR (middle) and a 1:1:1 mixture of Her2-Her2, Her2-Her3 and Her2-EGFR (bottom). NGS analyses were performed in duplicate and presented as mean values in the heatmap.

Next, we performed NanoDeep on Her2-expressing cancer cells. To verify the correlation between NGS reads from reference sequences and the amount of Her2-NanoCombs that were bound to the cells, we treated SKBR3 cells with increasing concentrations of Her2-NanoCombs. We found that there was a linear correlation between the levels of reference sequences and the concentration of NanoCombs. In subsequent experiments, NGS reads from reference sequences were used to scale the reads from detection sequences (Fig. 6a).

**Fig. 6:**
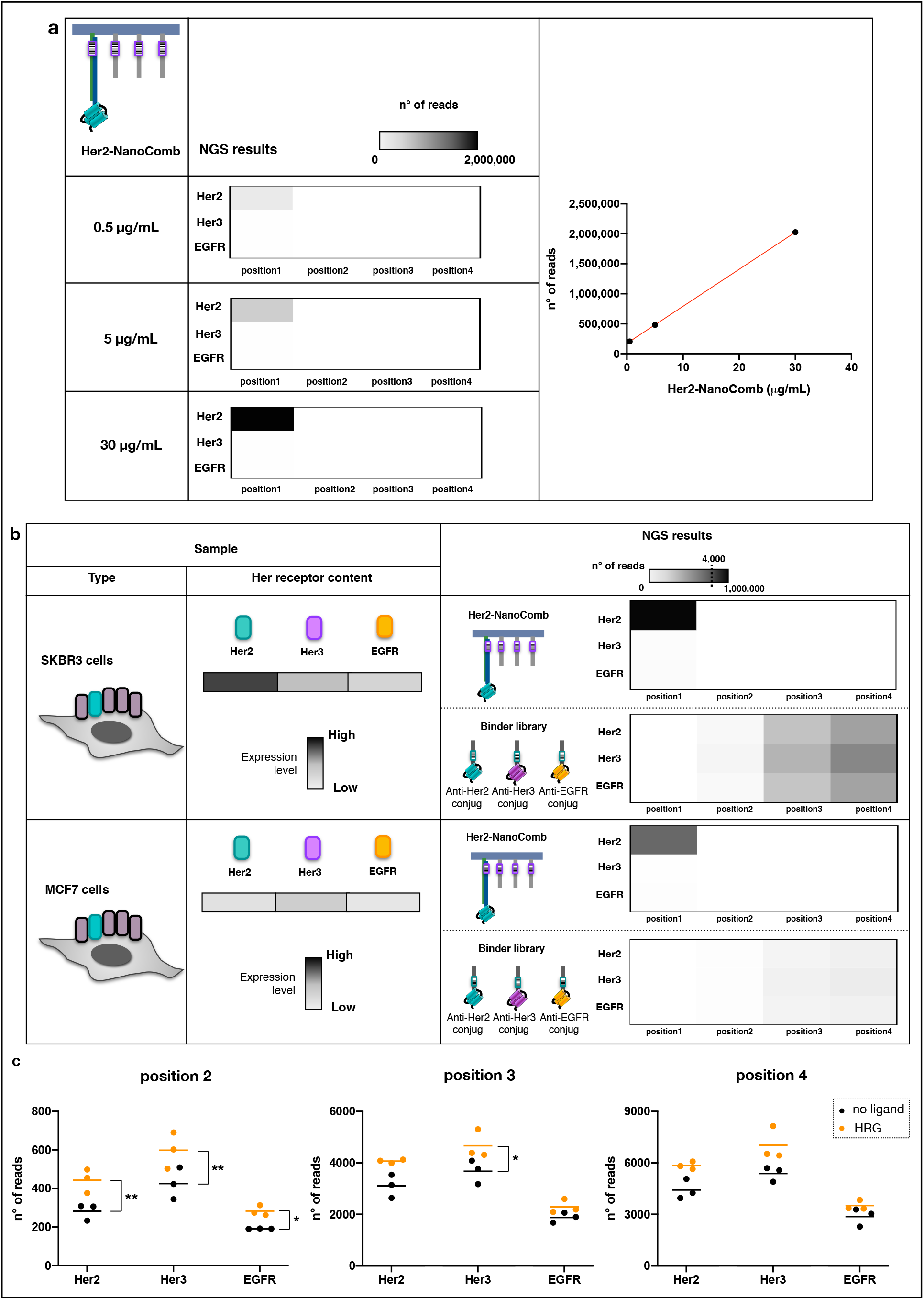
NanoDeep on cells. **a**, NGS analysis revealed a linear correlation between NGS reads and the concentration of Her2-NanoCombs. SKBR3 cells (400,000 cells/sample) were treated with three different concentrations of Her2-NanoCombs: 0.5 μg/ml, 5 μg/ml and 30 μg/ml. NGS reads are presented in heatmaps (left panel) and as a plot showing a linear correlation between the number of NGS reads of reference sequences and NanoComb concentrations (right panel). **b**, SKBR3 cells, Her2-overexpressing, and MCF7 cells, which show basal levels of expression of Her2, were grown to similar densities and analyzed by NanoDeep using Her2-NanoCombs and anti-Her2, -Her3 and -EGFR binder libraries. All measurements were performed in triplicate and presented as mean values in two types of heatmaps, showing the reads from the reference sequences (top) and detection sequences (bottom). **c**, SKBR3 cells were starved for 24 h and then stimulated for 15 min with HRG-β 1, followed by NanoDeep assay. Untreated cells were used as controls. For each position of the NanoComb, the number of reads of detection sequences was plotted versus the protein identity barcodes, for HRG-β1 treated cells (orange dots) and control cells (black dots). Dots represent three independent experiments and lines represent means of each condition. *p≤0.05; **p≤0.01.

We used NanoDeep to analyse Her2 nanoenvironments in SKBR3 and MCF7 breast cancer cells, which exhibit high and low expression levels of Her2, respectively. The rationale for testing cell lines with different expression levels is that the oncogenic capacity of Her2 is associated with the effects of overexpression on the frequency distributions of interactions between Her2 and other EGFR family members at the cell membrane^13–16^. NGS analysis of reference sequences demonstrated that the NanoDeep approach detected the differences in Her2 expression levels between the two cell lines. Analysis of the detection sequences provided information on the Her2 nanoenvironments. As shown in the heatmaps, the occupancies of Her2, Her3 and EGFR in Her2 nanoenvironments were similar in the two cell lines (Fig. 6b). However, the total levels of Her2, Her3 and EGFR in Her2 nanoenvironments were higher in SKBR3 cells compared to MCF7 cells. These results are consistent with previous reports showing that heterodimers between Her2 and other EGFR family members can form in Her2-expressing cancer cells, irrespective of Her2 expression levels^16, 19^. Together, these results support the notion that higher expression levels of Her2 lead to a higher number of interactions between Her2 and other EGF receptors resulting in increased oncogenic signalling in Her2-overexpressing cells compared to cancer cells with basal expression levels of Her2. Interestingly, the numbers of NGS reads consistently increased with increasing distance from the reference prong in NanoDeep experiments performed on cells (Fig. 6b). In contrast, on model SPR surfaces this effect was not observed.

To investigate whether NanoDeep can detect differences in cellular states due to ligand binding to membrane receptors, we treated SKBR3 cells with HRG-β1, one of the isoforms of Heregulin growth factor. HRG-β1 binds to Her3, inducing heterodimerization with Her2^30, 31^ and, to a lesser extent with EGFR^26^, leading to increased cell proliferation in breast cancer cell lines^16, 30, 32, 33^. NanoDeep revealed that treatment of SKBR3 cells with HRG-β1 led to increased occupancy of Her2 and Her3, and to a lesser extent EGFR, in the nanoenvironment of Her2 (Fig. 6c). The observed increase of Her3 in the nanoenvironment of Her2 is in line with the known mechanism by which HRG-β1 triggers the interaction between Her2 and Her3^16^. Further, the presence of increased levels of Her2 indicates that HRG-β1 binding to Her3 promotes membrane interaction networks that are more complex than pairwise interactions. Ligand binding to Her3 turns on the kinase activity of Her2 leading to oncogenic downstream signalling, a striking example of coupling between receptor dimerization and function. A molecular basis for the specific functions of higher-order structures has been described for EGFR, where self-association of ligand bound dimers leads to cooperativity in the activation of kinase domains ^23, 34^. However, the composition and roles of higher-order structures in EGFR family mediated signalling are still poorly characterized. NanoDeep has the potential to be a key tool in the investigation of the roles of higher-order association in EGFR family receptor signalling as well as in membrane receptor signalling in general.

## Conclusions

We developed a super-resolution method for unbiased mapping of protein nanoenvironments at the cell membrane, called NanoDeep. A simple DNA nanoassembly with encoded spatial information is used to decipher the spatial organization of proteins at the cell membrane, which are labelled with oligo-conjugated binders. NanoDeep combines multiplexed deciphering capability with nanoscale spatial resolution.

We applied NanoDeep to the analysis of the nanoenvironment surrounding the membrane protein Her2. We found that the composition of Her2 nanoenvironments did not differ significantly in two cancer cell lines with vastly different levels of Her2 expression. However, cell treatment with HRG-β1, a ligand for Her3, led to increased recruitment of Her3 and Her2 to Her2 nanoenvironments.

NanoDeep has the potential to provide a breakthrough in the simultaneous analysis of the spatial distribution of many proteins at the membrane without microscopy measurements. The ability to obtain ensemble averaged data, that is free from the sampling and confirmation bias of microscopy, will provide a powerful tool for understanding the importance of membrane protein assemblies.

## Methods

### DNA sequences

All synthetic DNA sequences were obtained from Integrated DNA Technologies. DNA sequence details are reported in the Supplementary Information.

### Native PAGE and Electro Mobility Shift Assay (EMSA)

Native PAGE gels were prepared as follows: a fresh solution comprising of polyacrylamide (19:1) at final concentrations from 7 to 13% (Biorad), 0.5x TBE buffer (45 mM Tris-borate, 1 mM EDTA) and glycerol (2.5%, v/v) were mixed and polymerized by the addition of fresh 10% APS solution (Merck Millipore) (final concentration 10%) and TEMED (Biorad) (final concentration 1%) in a mini-protean gel system (Biorad). TBE 0.5× was used as running buffer and the gels were run at room temperature for 40 min at 200 V. Depending on the different applications, gels were stained for nucleic acids with SybrGold (Thermofisher Scientific) and for total protein with GelCode blue stain reagent (Thermofisher Scientific).

### Folding, purification and electrophoretic characterization of DNA NanoCombs

Backbone and protruding strands (prongs) ssDNA sequences were diluted at a concentration of 1 μM and 2 μM, respectively, in Hybridization Buffer (5 mM Tris-HCl pH 8.0, 10 mM MgCl_2_, 1 mM EDTA). Folding was carried out by rapid heat denaturation (80 °C for 10 min) followed by cooling to RT for 2 h. Removal of excess prongs was done first with washing in 50 kDa MWCO 0.5 mL Amicon centrifugal filters (Merck Millipore) and then by means of Streptavidin-coated magnetic beads (DYNAL MyOne Dynabeads Streptavidin C1-Thermofisher Scientific) as follows: beads were washed with PBS and then with Immobilization Buffer (5 mM Tris-HCl pH 7.5, 1 M NaCl, 0.5 mM EDTA). NanoCombs were incubated with beads at RT for 1 h; beads were then collected with a magnet and the supernatant was discarded. The elution of NanoCombs from beads was performed with 5 mM biotin (Invitrogen) for 30 min at RT. Native polyacrylamide gels were stained with SybrGold to visualize the NanoComb stepwise assembly (7%) and the purification yield (13%).

### Design, expression and purification of affibody-VirD2 fusion proteins

The coding sequences of anti-Her2 (Z_Her2:342_^35^), anti-Her3 (Z_08699_^36^) and anti-EGFR (Z_EGFR:2377_^37^) affibodies were synthetized (BioCat) and cloned (XhoI/BamHI) at the C-terminal of the VirD2 protein connected via a flexible linker (GGGGS) in the expression plasmid pET-16-b and the sequences were validated by sequencing. Expression and purification of recombinant proteins were carried out as previously described^27^ at the Karolinska Institutet/SciLifeLab Protein Science Core Facility.

### VirD2 tag mediated conjugation of oligonucleotides to affibodies and purification of conjugates

For the conjugation reaction, protein and oligonucleotide were mixed at a 1:0.75 ratio in TKM buffer (50 mM Tris-HCl pH 8, 150 mM KCl, 1 mM MgCl_2_, 10% glycerol) and incubated at 37 °C for 2 h. Conjugates were then purified from the excess oligo and the fraction of protein not conjugated by isolation from a native PAGE gel. Briefly, conjugates were separated from oligos and excess protein by electrophoresis by native PAGE (10%). The bands corresponding to the conjugates were cut and incubated overnight at 4 °C in TBS buffer (50 mM Tris-HCl pH 7.5, 150 mM MgCl_2_). Buffer with eluted conjugates was finally filtrated with Nylon 0.45 μM centrifugal filter (Thermofisher Scientific).

### Functionalization of NanoCombs with anti-Her2 affibody-oligo conjugates on reference prong (Her2-NanoCombs)

NanoCombs were folded and loaded on Streptavidin-coated magnetic beads, as previously described. Anti-Her2 affibody, conjugated with a DNA sequence that partially hybridizes to the reference prong and purified as described above, was incubated with the beads at RT for 2 h. Beads were then collected with a magnet and the supernatant was discarded. Finally, Her2-NanoCombs were eluted from the beads with 5 mM biotin for 30 min at RT.

### Surface Plasmon Resonance assays

Biacore T200 instrument and related reagents (GE Healthcare) were used to perform all Surface Plasmon Resonance experiments. *<u>NanoComb assembly characterization</u>*. SA gold sensor chip was used to immobilize desthiobiotinylated-backbone DNA sequence (ligand) and to verify the hybridization of the protruding strands (analyte). A continuous flow of running buffer HBS-EP+ buffer (20 μL/min) was maintained during both ligand immobilization and analyte injections. *<u>Binding/kinetics of VirD2-affibodies and affibody-olgo conjugates</u>*. ECD-Her2/-Her3/-EGFR proteins (AcroBiosystem), diluted at 10 μg/mL in 10 mM Na acetate, pH 4.5, were immobilized on CM3 or CM5 gold sensor chip at the desired surface densities via amine coupling reactions, according to the manufactures’ instructions. Binding affinity tests of the VirD2-affibodies and affibody-oligo conjugates were performed by injecting different concentrations of analytes in running buffer at 50 μL/min. For the regeneration of the surface, 18 s pulses of a Gly-HCl pH 2.0 solution were used, followed by a stabilization time of 5 min. Target selectivity tests were performed by injecting a single concentration (50 nM) of affibody-oligo conjugates over the three targets immobilized separately on three different flow cells of the same sensor chip. HBS-EP+ buffer was flowed as running buffer over the surface at 5 μL/min for protein immobilization and at 50 μL/min for binding analysis. The dissociation equilibrium constant (KD), the association rate constant (kon), and the dissociation rate constant (koff) were determined using the BIAevaluation 3.0 software, assuming a 1:1 binding model. *<u>Characlerizalif/n of the binding between conjugates and NanoCombs</u>*. Desthiobiotinylated-NanoCombs were immobilized on SA gold sensor chips. Then affibody-oligo conjugates (targeting Her2) were injected over the surface at 1 μM. The binding capability of Her2 was verified by flowing three increasing concentrations of ECD-Her2 (0.5-5-50 nM) in single cycle kinetic mode. In the reverse assay, first ECD-Her2 was covalently immobilized, as previously described, on CM3 sensor chips, reaching three different surface densities on the three independent flow cells. Next, anti-Her2 affibody-oligo conjugates (at 50 nM) and NanoCombs (at 400 nM) were sequentially flowed over the surface. HBS-EP+ buffer was used as running buffer (flow rate of 5 μL/min for immobilization and at 30 μL/min for binding assays). *<u>Toehold exchange strategy</u>*. Assay 1: Binder oligos corresponding to the DNA sequences of the affibody-oligo conjugates, biotinylated at the 5’ end, were immobilized on SA sensor chips. Blocking strand oligos were injected at the concentration of 1 μM; the displacement of blocking oligo by branch migration was promoted by injection of 2 μM of invading strands over the surface. After that, a partially assembled NanoComb (backbone + prong 2 and prong 3) at 2 μM concentration was allowed to interact with the unblocked immobilized oligos.

TE/Mg^2+^ buffer (10 mM Tris pH 8.0, 12.5 mM MgCl_2_,1 mM EDTA) was used as running buffer (flow rate of 20 μL/min both for immobilization and binding assays). Assay 2: 10 μg/mL of ECD-Her2 and ECD-Her3 were mixed 1:1 and incubated for 1 h. Then they were injected over a CM5 sensor surface (two flow cells) for covalent immobilization, as previously described. Binding of Her2-NanoCombs was performed by multiple injections to obtain a binding level over the surface of ~250 RU. After that, anti-Her3 affibody-oligo conjugates, previously hybridized with blocking strand, were injected at 5 μM over only one of the two active flow cells. The blocking strand was then displaced by injection of invading strand 1 at 10 μM; during this step the temperature was increased at 45 °C to prevent hybridization of invading strand to the detection prongs. Temperature was decreased at 25 °C and the system was left in stand-by mode (flow rate of running buffer at 5 μL/min) for 1 h to allow hybridization of conjugates oligo to the detection prongs. After that, invading strand 2, specific for the toehold-mediated displacement of anti-Her2 conjugate from the reference prong, was injected at 4 μM. HBS-EP+ buffer was used as running buffer (flow rate of 5 μL/min for immobilization and at 10-30 μL/min for binding/displacement steps). *<u>Validation of enzymatic reaclions</u>*. ECD-Her3 was covalently immobilized, as previously described, on two flow cells of CM5 sensor chip, reaching the same immobilization level. Anti-Her3 affibody-oligo conjugates were injected over the surface at 500 nM, followed by injection of NanoCombs at 1 μM in HBS-EP+ as running buffer. After that, T4 polymerase (New England BioLabs) at 15 units/mL was injected over the flow cell 1 in Neb2.1 buffer 1x (New England BioLabs) with a flow rate of 5 μL/min for 15 min at 12 °C. Then, first EcoRI (New England BioLabs) and afterwards BamHI (New England BioLabs), at 15 units/mL, were injected over the flow cell 1 in Neb2.1 buffer 1x and Cut Smart buffer 1x (New England BioLabs) respectively, with a flow rate of 5 μL/min for 15 min at 37 °C. Finally, running buffer was changed back to HBS-EP+ to perform the injection of Streptavidin at 1 μM over both flow cells 1 and 2. *<u>NanoDeep experiment</u>*. Different sensor surfaces were created by covalently immobilizing ECD-Her2/-Her3/-EGFR alone or in different combinations on CM5 chip. Her2-NanoCombs were injected over the surface. After that, libraries of affiboby-oligo conjugates, previously hybridized with blocking strand, were injected at 5 μM. The temperature was increased to 45 °C and the blocking strand was then displaced by injection of invading strand at 10 μM. Temperature was decreased to 25 °C and the system was left in stand-by mode (flow rate of running buffer at 5 μL/min) for 1 h to allow hybridization of binder oligos to the detection prongs. Enzymatic reactions by T4 polymerase, EcoRI and BamHI were performed as previously described. As the final step, the digestion products of BamHI were recovered in a 2 mM EDTA solution to stop the reaction.

### Cell culture and cell samples preparation

Human cancer cell lines SKBR3 and MCF7 (American Type Culture Collection) were cultured in 5% CO_2_ at 37 °C in DMEM (Gibco) supplemented with 10% fetal bovine serum (Thermofisher Scientific) and 1% penicillin–streptomycin (Thermofisher Scientific). To perform the NanoDeep protocol, cells were plated on 35 mm-diameter culture dishes and allowed to adhere for 24 h. Then cells were fixed with 4% formaldehyde for 20 min and washed with PBS. For ligand treatment assays, SKBR3 cells were starved for 24 h in serum-free medium and then stimulated for 15 min with 7-15 nM HRG-β1 (Sigma Aldrich).

### NanoDeep on cells

Fixed SKBR3 and MCF7 cells, were incubated for 40 min at RT with blocking buffer (50 mM Tris-HCl pH 7.5, 150 mM NaCl, 5 mM EDTA, 250 μg/mL BSA, 15 μg/mL salmon sperm DNA (Invitrogen)) to prevent non-specific binding (Fig. S5). Three times washing with PBS + 0.05% Tween20 was then performed and, after that, Her2-NanoCombs were diluted in PBS + BSA 3% and incubated for 2 h at RT, followed by washing. Affibody-oligo conjugate libraries, previously hybridized with blocking strand, were next diluted in PBS + BSA 3% and added over the cells, followed by washing. Cells were then incubated at 45 °C and, after reaching the temperature, the displacement of blocking strand was performed by addition of invading strand (1 h), followed by washing. After that, cells were left at RT for 3 h to allow hybridization of binder oligos to the detection prongs. Enzymatic reactions by T4 polymerase, EcoRI and BamHI were performed as previously described. Finally, the digestion products of BamHI were recovered and supplemented with 2 mM EDTA to stop the reaction, and then concentrated by means of 3 kDa MWCO 0.5 mL Amicon centrifugal filters.

### PCR amplification

GoTaq Hot start polymerase (Promega) was used for amplification of barcoded sequences derived from NanoDeep experiments. PCR was performed with the following conditions: 95 °C for 2 min, 30 cycles of 94 °C for 30 s, 38 °C for 30 s and 72 °C for 3 s, and then 72 °C for 5 min. M13F sequence (GTTTTCCCAGTCACGAC) was added at 5’ of 17-nt primers to reduce non-specific PCR products.

### NGS sequencing

Sequencing libraries were prepared using the ThruPLEX Tag-seq Kit (Takara Bio). 10 μL of samples from NanoDeep assays (performed on SPR surfaces or cells) were processed by the three-steps workflow described in the kit protocol. Reaction products were purified with Ampure-XP beads (Beckman Coulter) as follows: 0.9x volume of beads were added to 50 μL of samples and incubated for 10 min. Beads were collected with a magnet and the supernatant was transferred to different tubes and incubated with 1.8x volume of new beads for 10 min. After washing with 80% EtOH (30 s, two times), DNA was eluted with 20 μL of EB buffer (Qiagen).

Sequencing was performed using the NextSeq 550 instrument (Illumina), following the manufacturer’s standard protocol. The samples were single-end sequenced with a read length of 75 bp and 8 bp index reads.

### NGS Sequencing Data Analysis

The resulting sequences were analyzed through cataloguing molecules by UMI’s and identifying experimental barcodes using custom code written in Python and utilizing functions from the Numpy and Biopython libraries. Sequences were first processed to identify and catalog each read by its UMI according to the Tagseq pipeline. Sequences were then analyzed for presence of barcode concatenation by searching by pairwise alignment (Smith-Waterman) for a common sequence and its reverse complement sequence which would be flanked by two 6-nt barcode sequences from the set of known protein identities and associated positions in the event of a concatenation. A threshold score of 80% match with the common sequence was used to select candidates for barcodes identification. Subsequent candidates were further filtered according to whether an 80% match with one of the known barcodes could be found in the 6 nucleotide positions immediately 5’ upstream and 3’ downstream of the common sequence. Incidence was then tallied for each pair of barcodes identified and compiled into association matrices. Each tally was weighted by dividing its value by the UMI incidence for that particular read in order to eliminate amplification bias among associations. Code for UMI processing and barcode association are available online at https://github.com/xx.

### Statistical analysis

Statistical analysis was carried out with GraphPad Prism (Version 8.2.1). Statistical significance was determined by performing two-tailed Student’s *t*-test. A *p*-value ≤ 0.05 was considered statistically significant.

## Data availability

All data supporting the results of this study have been included in the main text and supplementary information. Raw data are available from the authors on reasonable request.

## Acknowledgements

The authors acknowledge Björn Reinius for helpful discussions. A.I.T. acknowledges support from the European Research Council under the European Union’s Seventh Framework Programme (ERC, grant no. 617711), the Swedish Research Council (grant no. 2015-03520) and the Knut and Alice Wallenberg Foundation (grant no. KAW 2017.0114).

## Authors contributions

E.A. designed the study and performed the experiments; G.B. and B.H. provided key insights for the design of experiments; I.H developed NGS data analysis; L.H and R.S. contributed to performance and interpretation of NGS experiments; A.I.T. conceived and supervised the study; E.A. and A.I.T. wrote the manuscript, with input from all authors; all authors contributed to the manuscript revision and gave approval to the final version.

## Competing interests

The authors declare no competing interests.

## Corresponding author

Correspondence to Ana I. Teixeira.

