## Supplemental information for "A DNA nanoassembly-based approach to map membrane protein nanoenvironments"

### Supplementary information

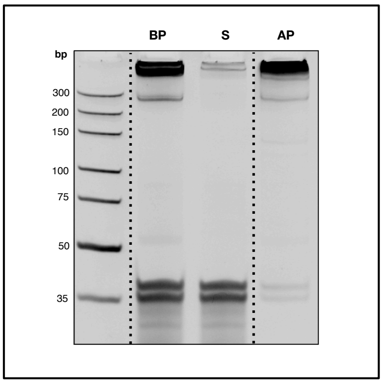


**Fig. S1: NanoComb purification.** NanoCombs were purified from excess of prongs that were not hybridized to the backbone using Streptavidin (SA)-coated magnetic beads. NanoCombs were bound to beads by means of a desthiobiotin group present at the 3' end of backbone and were collected using a magnet. The release of NanoCombs from the beads was then performed by biotin elution. Native PAGE (13%) was used to analyze the purification of NanoComb samples. BP, before purification; S, discarded supernatant; AP, after purification.

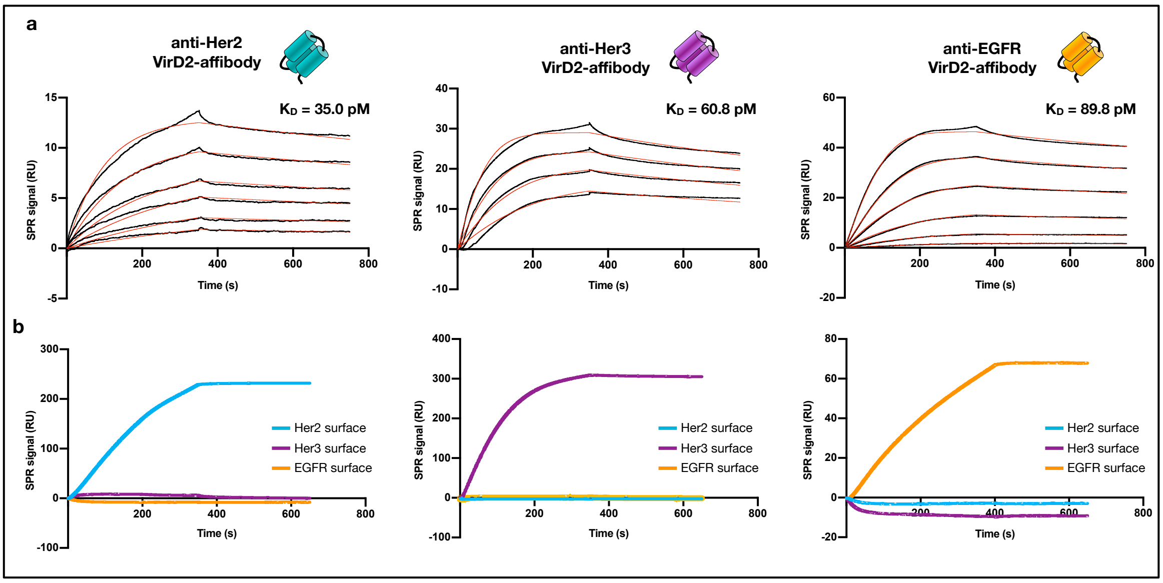


**Fig. S2: Binding affinity and target selectivity characterization of VirD2-affibody fusion proteins. a,** SPR binding analysis of VirD2-affibody fusion proteins targeting Her2, Her3 and EGFR performed at concentrations ranging between 0.33 nM and 5 nM, to cover the kinetic spectrum. Fitting was performed using a 1:1 kinetic model to determine the dissociation constant, $K_D$. Recorded sensorgrams are shown in black, while fitting curves are in red. **b,** Selective binding of VirD2-affibody fusion proteins to their specific targets was verified by recoding the binding sensorgrams of VirD2-affibodies to the ECDs of Her2, Her3 and EGFR immobilized on three different flow cells.

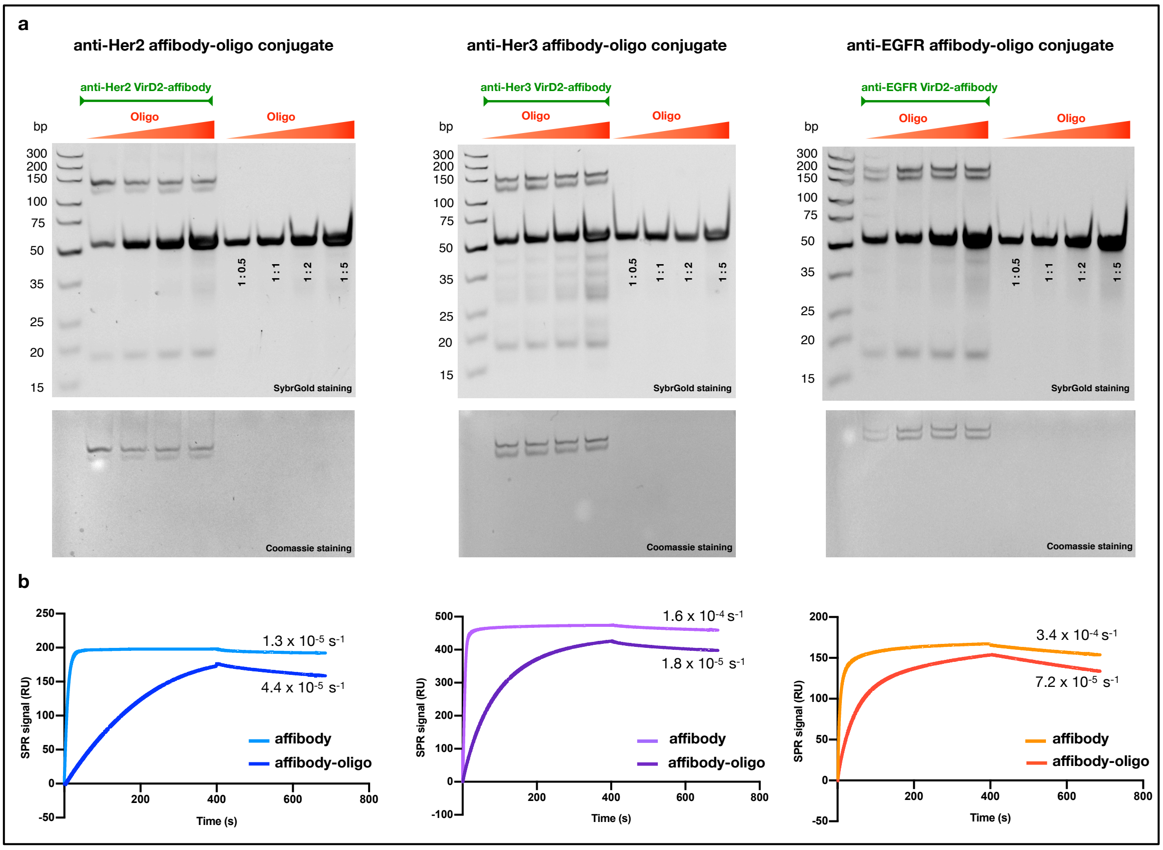


**Fig. S3: Characterization of affibody-oligo conjugates. a,** Equivalent amounts of anti-Her2-, anti-Her3- or anti-EGFR-VirD2-affibody fusion proteins were incubated with increasing concentrations of their corresponding oligos for 2 h at 37 °C. Native PAGE (10%) was used to detect DNA by staining with SybrGold. The same gel was stained with Coomassie Blue to visualize proteins, thereby revealing the formation of DNA-protein conjugates. **b,** SPR real-time kinetic analysis of VirD2-affibodies (light color) and the respective VirD2-affibody-oligo conjugates (dark color) on sensor surfaces functionalized with their respective target proteins. The dissociation constants ($k_{off}$) reported for each sensorgram were determined by fitting with 1:1 kinetic model.

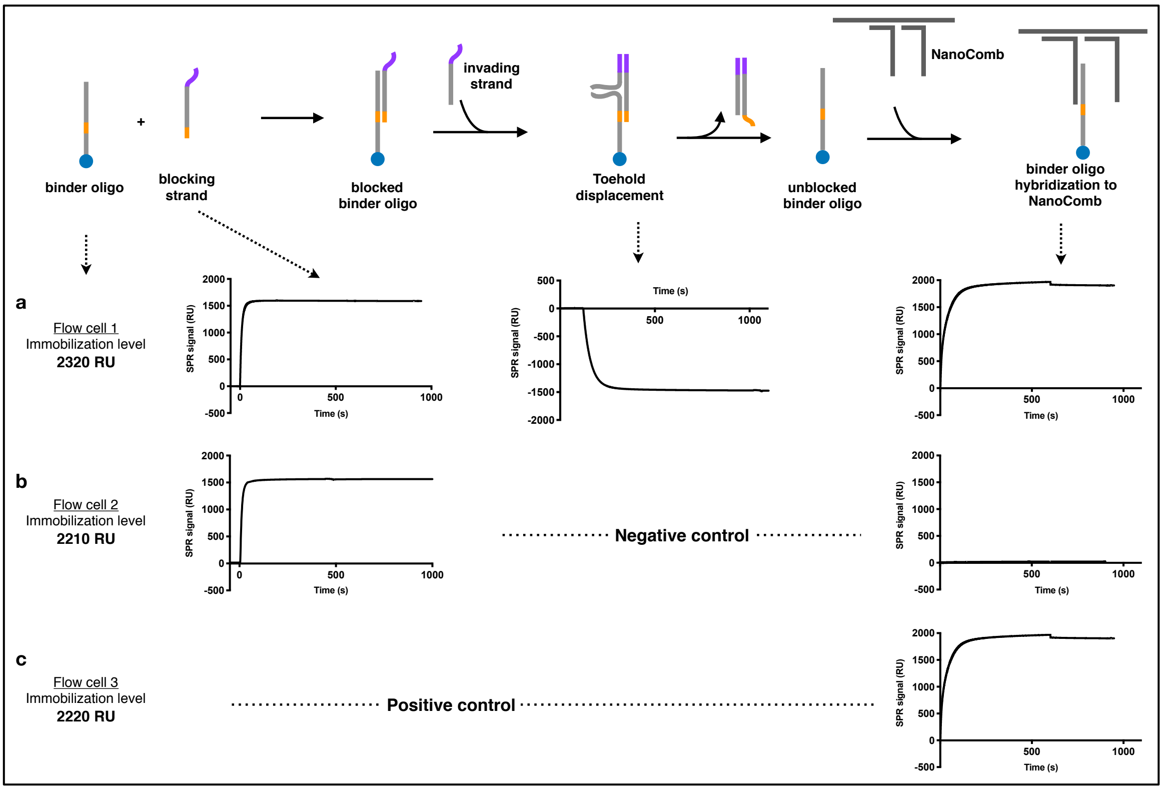


**Fig. S4: Toehold exchange reversibly blocked the hybridization of oligos to the NanoCombs.** Biotinylated versions of the oligos used to produce the affibody-oligo conjugates were immobilized onto SA sensor surfaces. **a,** Hybridization with the blocking strand caused an increase in the SPR signal, which was followed by a decrease in the signal when the invading strand was injected. The resulting unblocked oligos were capable of hybridizing to the NanoCombs. **b,** Without the strand invasion step, the blocked oligos were not able to bind to the NanoCombs (negative control). **c,** In the positive control there were no blocking/unblocking steps and the NanoComb hybridized to the anchored oligos.

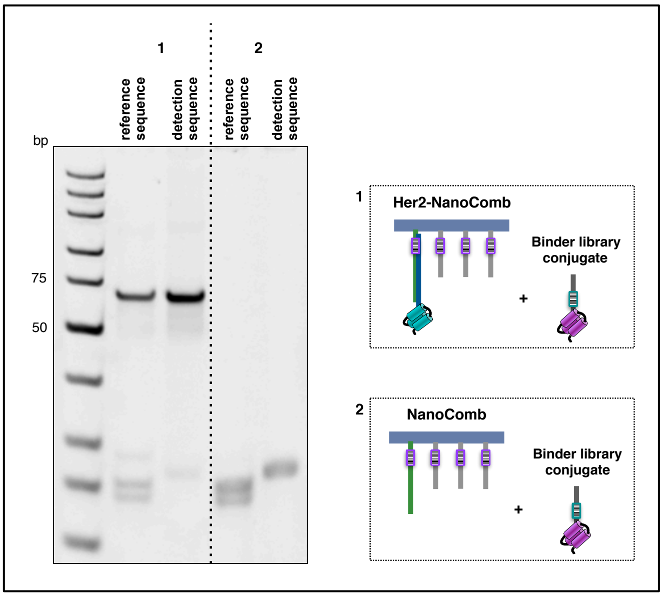


**Fig. S5: Assessment of the specificity of the binding of NanoCombs to cells.** NanoDeep was performed on SKBR3 cells, using Her2-NanoCombs or NanoCombs that were not functionalized with anti-Her2 affibody-oligo conjugate on the reference prong, as a negative control. Reference and detection sequences, amplified by PCR and visualized on native PAGE (13%), were recovered only in presence of Her2-NanoComb (1) and not on the negative control (2), showing that NanoCombs bound to the cells through a specific interaction between the binder and the reference protein and not through a non-specific DNA interaction with the cell surface.

### Supplementary Table 1: DNA sequences

| Name | Sequence |
|---|---|
| **Oligos for NanoComb** | |
| Backbone | G A A T C G G T T C T A G C A T T C G C A T A C A A A C T C T G G A G A G G C A A A G A C A C G T T A C A G A G A<br>A A A C A T C T A C C A C A G G A G G A G A T C C C T C A C C G A G C C T A G A A G G -desthiobiotin |
| prong1 | C C T T C T A G G C T C G G T G A G G G A T T T T G A A T T C **A A A C T G** A G G G A A A G A A A T C A G A G C T T |
| prong1 _for "bare" NanoComb | C C T T C T A G G C T C G G T G A G G G A T T T T G A A T T C **A A A C T G** G G T A T T G A A T G A G T G |
| prong2 | T C T C C T C C T G T G G T A G A T G T T T T T T G A A T T C **C G G C G T** G G T A T T G A A T G A G T G |
| prong3 | T T C T C T G T A A C G T G T C T T T G C T T T T G A A T T C **A T A G G C** G G T A T T G A A T G A G T G |
| prong4 | C T C T C C A G A G T T T G T A T G C G A T T T T G A A T T C **A C G T A C** G G T A T T G A A T G A G T G |
| end | A T G C T A G A A C C G A T T C |
| **Oligos for conjugates** | |
| AbCodeHer2_NanoComb_p1 | C T C A A A T T A C A A C G G T A T A T A T C* C* T* G* C* C* A* G T A G T T T T G G A T C C **A G A G T C** A A G C T C T G<br>A T T T C T T T C C C T |
| AbCodeHer2_NC_p1_toehold displ. | C T C A A A T T A C A A C G G T A T A T A T C* C* T* G* C* C* A* G T A G T T T T G G A T C C **A G A G T C** A A G C T C T G<br>A T T T C T T T C C C T G G A C A T A |
| AbCodeHer2_library | C T C A A A T T A C A A C G G T A T A T A T C* C* T* G* C* C* A* G T A G T T T T G G A T C C **A G A G T C** G C A T C C A C<br>T C A T T C A A T A C C |
| AbCodeHer3_library | C T C A A A T T A C A A C G G T A T A T A T C* C* T* G* C* C* A* G T A G T T T T G G A T C C **A A C A G T** G C A T C C A C<br>T C A T T C A A T A C C |
| AbCodeEGFR_library | C T C A A A T T A C A A C G G T A T A T A T C* C* T* G* C* C* A* G T A G T T T T G G A T C C **C A G A C T** G C A T C C A C<br>T C A T T C A A T A C C |
| AbCode_library_toehold displ. | Biotin - T T T T G G A T C C A A C A G T G C A T C C A C T C A T T C A A T A C C |
| **Oligos for toehold displacement** | |
| Blocking strand | A G T C G A G G T A T T G A A T G A G T G G A T G C |
| Invading strand (1) | C A C T C A T T C A A T A C C T C G A C T |
| Invading strand 2 | T A T G T C C A G G G A A A G A A A T C A G |
| **PCR primers** | |
| Primer_Ab_reference seq | G T T T T C C C A G T C A C G A C C A G A G T C A A G C T C T G A T |
| Primer_Ab_detection seq | G T T T T C C C A G T C A C G A C C A G A G T C G C A T C C A C T C |
| Primer_p1 | G T T T T C C C A G T C A C G A C C A A A C T G A G G G A A A G A A |
| Primer_p2 | G T T T T C C C A G T C A C G A C C C G G C G T G G T A T T G A A T |
| Primer_p3 | G T T T T C C C A G T C A C G A C C A T A G G C G G T A T T G A A T |

*phosphorothioate bonds is introduced in the conjugation site to further increment the reaction yield[1].
Bases in bold correspond to barcode sequences.

1. Bernardinelli, G. & Högberg, B. Entirely enzymatic nanofabrication of DNA-protein conjugates. *Nucleic Acids Res* **45**, e160 (2017).
